# Interleukin-22 in enteroendocrine cells controls early-life gut motility through interactions with the microbiota

**DOI:** 10.1101/2025.04.25.650580

**Authors:** Soraya Rabahi, Emiliano Marachlian, Fabian Guendel, Aya Mikdache, Keinis Quintero-Castillo, Lucie Maurin, Vincenzo Di Donato, Akshai Janardhana Kurup, Yazan Salloum, Gwendoline Gros, Patricia Diabangouaya, Camila Garcia-Baudino, Ignacio Medina-Yáñez, Jean-Pierre Levraud, Georges Lutfalla, Filippo Del Bene, Jos Boekhorst, Carmen G. Feijoo, Gerard Eberl, Sylvia Brugman, German Sumbre, Pedro P. Hernandez

## Abstract

The gut microbiota, immune system, and enteric nervous system tightly interact to regulate adult gut physiology. Yet the mechanisms establishing gut physiology during development remain unknown. Here, we report that in developing zebrafish, enteroendocrine cells produce IL-22 in response to microbial signals before lymphocytes populate the gut. In larvae, IL-22 is crucial to set gut microbiota composition, and ghrelin hormone expression to promote gut motility. IL-22 developmental function depends on its ability to modulate gut microbiota, as bacterial transfer from wild-type zebrafish restored gut motility in *il22*^-/-^ by reestablishing ghrelin hormone expression. Additionally, IL-22-deficient mice show impaired gut motility and reduced ghrelin expression in early life, indicating a conserved function. Altogether, we identify a circuit where enteroendocrine cells regulate gut function via cytokine control of the microbiota, showing how gut physiology is set prior to immune system maturation.

## Introduction

The mature gut integrates and controls signaling across the immune, nervous, and endocrine systems to regulate food intake and digestion. In the adult, these functions are achieved through the precise orchestration of diverse cell types from each of these systems. During postembryonic, not all of these cell types are present, yet how intestinal physiology is ensured remains unknown. In particular, this developmental stage is marked by exposure to the external environment and colonization by complex microbiota, critical for the maturation of intestinal cells, including epithelial, neuronal, and immune cells (*1–4*). In zebrafish, although mature lymphocytes appear at 2-3 weeks of age (*5*), ingestion starts as early as four days post-fertilization. How intestinal physiology and homeostasis are achieved in developing animals, which lack a mature immune system, remains unknown.

Gut lymphocytes produce cytokines with key roles in adult gut homeostasis. Among them, interleukin-22 (IL-22) is particularly important since its receptor is mainly expressed in epithelial cells (*6*), while IL-22 itself is expressed in immune cells such as T cells and innate lymphoid cells (ILCs) (*6, 7*). Studies in mice have shown that IL-22 regulates gut barrier integrity (*8–10*), tissue repair (*11*), host metabolism (*12*), and microbiota composition (*13*). However, whether and how cytokines impact both the microbiota and gut physiology during early-life, as well as their cellular sources, remains poorly understood.

While lymphocytes become abundant in the gut only at 2-3 weeks of age in both mice and zebrafish, epithelial enteroendocrine cells (EECs) are present and functional before this age (*14, 15*). EECs are conserved across insects, fish, and mammals, and orchestrate gut physiology, including gut motility, by secreting hormones and neurotransmitters in response to diverse stimuli (*16–18*). The microbiota is able to exert its function in large part by stimulating the release of EEC signaling molecules (*14, 19*). EECs thus are a prime candidate for regulating both early gut function and interactions with the microbiota during development.

Here, we identified an essential function for IL-22-microbiota interactions in early-life gut physiology. We show that tryptophan metabolites induced IL-22 expression in EECs through activation of the transient receptor potential ankyrin 1 (Trpa1b) channel in the larval zebrafish gut, prior to immune system maturation. Furthermore, IL-22 deficiency not only resulted in defects in microbiota composition and increased susceptibility to gut bacterial infection but also impaired gut neuronal activity and gut motility. The gut motility impairment in *il22*^-/-^ was rescued by microbiota transfer from WT larvae. Transcriptomic analysis revealed that the intestines of *il22*^-/-^ larvae have dysregulated expression of several EEC-derived hormones. Microbiota transfer from WT larvae restored expression of the hormone *ghrelin*. Furthermore, incubation of *il22*^-/-^ larvae with ghrelin peptide restituted gut motility, while blocking ghrelin activity impeded the rescue promoted by co-housing. Analysis of *Il22*^-/-^ mice revealed impaired food transit and reduced ghrelin expression in the gut during early life, suggesting a conserved role for IL-22 in maintaining gut physiology in developing animals. Together, these results reveal novel mechanisms by which host-microbiota interactions regulate gut function during development.

## Results

### *il22* expression is induced in enteroendocrine cells upon stimulation from bacterial cues

To determine whether and how IL-22 plays a role in the developing gut, we first analyzed its expression in zebrafish larvae. Given that zebrafish larvae have only a few lymphocytes, we aimed to identify the cell types expressing this cytokine in their gut. To determine whether IL-22 is expressed in intestinal epithelial cells (IECs)—the predominant cell type in the gut— we generated a *il22*:mCherry reporter line and crossed it with the *cldn15la*:GFP line (*20*), which specifically labels IECs. We observed *il22*:mCherry expression in scattered IECs (**Fig. 1A)**. Further analysis of scRNA-seq data from larval intestines (*21*) confirmed that IECs, particularly EECs, were the primary source of *il22* (**Fig. S1A, B**). Imaging *il22*:mCherry crossed with the *neurod1*:GFP line (*22*), which specifically labels EECs in the gut, showed co-localization, with 90,5% ± 6,2 (n=7 larvae) mCherry^+^ cells being also GFP^+^ (**Fig. 1B, Table S2**). Our results identify enteroendocrine cells as the major source of *il22* in the zebrafish larval gut.

**Figure 1.**
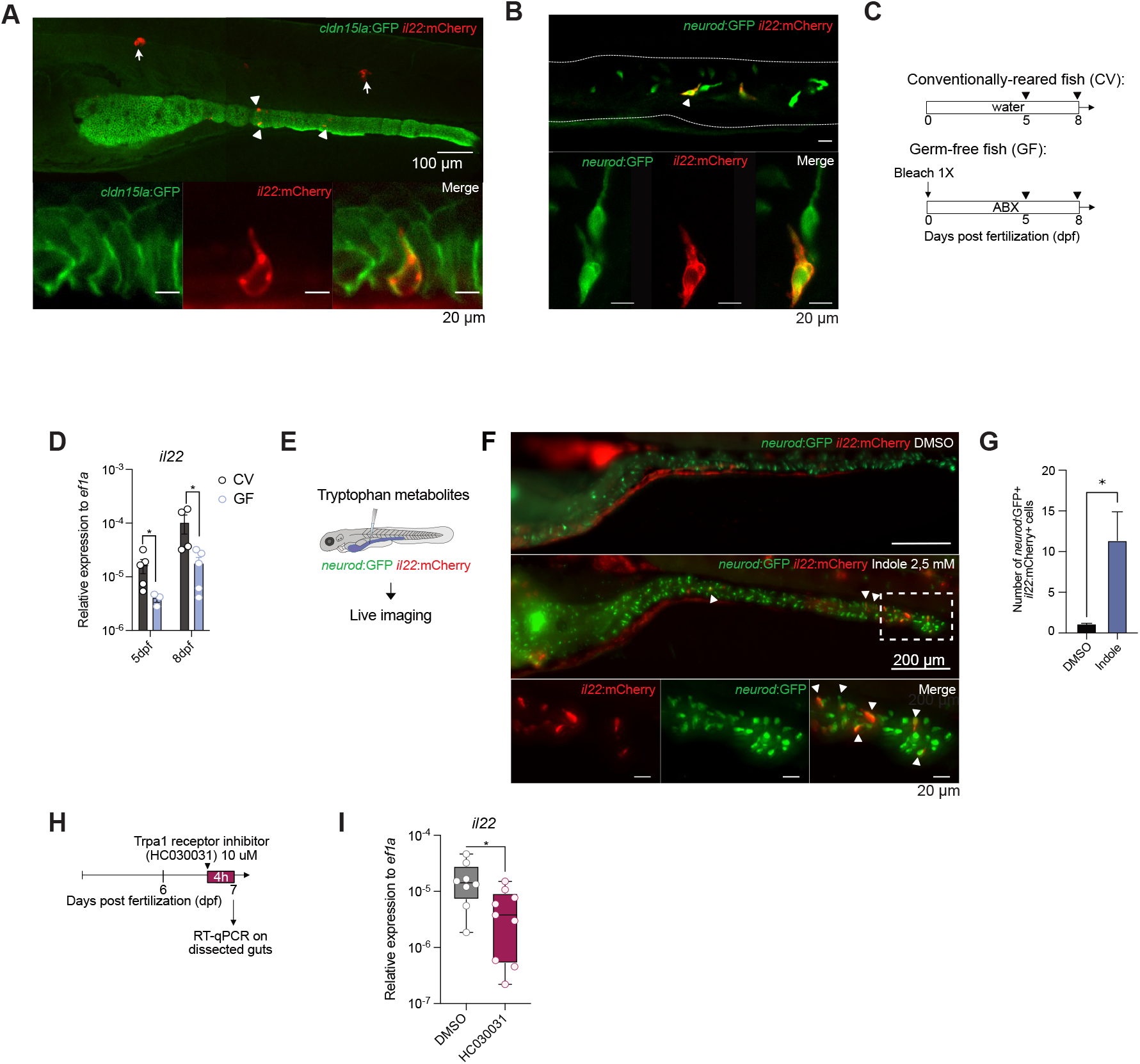
*il22* expression is induced in enteroendocrine cells upon bacterial cues. **A.** Confocal images of 7 dpf Tg(*cldn15la*:GFP, *il22*:mCherry) larvae. Arrows show neuromasts and arrowheads show IECs expressing *il22*. Scale bar = 100 um (top panel) and 20 um (bottom panel). **B**. Confocal images of Tg(*neurod*:GFP, *il22*:mCherry). Arrows show co-localization of the two transgenic markers. Scale bar: 10 um. **C**. Diagram showing bleach and antibiotic (Abx) treatment in zebrafish embryos/larvae to generate germ-free (GF) animals. Black arrows indicate the age of the larvae when guts where dissected. **D**. RT-qPCR analysis of *il22* expression in dissected guts from 5 and 8 dpf conventionally reared (CV) or GF larvae. **E**. Schematic representation of the tryptophan metabolite indole injection in the guts of 7 dpf Tg(*neurod*:GFP, *il22*:mCherry) larvae, followed by live imaging 4 hours later. **F**. Fluorescent images of Tg(*neurod*:GFP, *il22*:mCherry) injected with PBS (top) or indole (2,5 mM) (middle). Zoom-in view (bottom) shows *neurod*:GFP^+^, *il22*:mCherry^+^ cells in the posterior intestine. Arrowheads indicates double positive cells. Scale bar: 200 um (top and middle), 20um (bottom). **G**. Quantification of the number of *neurod*:GFP+ *il22*:mCherry+ cells in the gut of 7 dpf larvae injected with PBS or indole. **H**. Diagram showing the treatment of 6 dpf larvae with the Trpa1 inhibitor HC030031 (10 uM) during 4 hours before dissecting guts a 7 dpf. **I**. RT-qPCR on dissected guts after treatment with the Trpa1 inhibitor HC030031 or DMSO. Statistical analysis was performed with Mann-Whitney * *P* < 0,05. Data are representative of at least two independent experiments.

Previous studies in mammals revealed a central role of the microbiota in inducing IL-22 expression in the mature gut (*23*). We compared germ-free (GF) to conventionally reared (CV) zebrafish larvae to determine whether the microbiota could influence *il22* expression level in EECs. GF larvae had lower gut *il22* mRNA levels compared to CV counterparts, indicating that the microbiota promotes baseline *il22* expression (**Fig. 1C, D**). We then aimed to identify the mechanism by which the microbiota influences *il22* expression in EECs. In mammals, dietary and bacteria-derived tryptophan metabolites trigger IL-22 production in lymphocytes *via* the aryl hydrocarbon receptor (AhR) (*24*). The *il22*-expressing EECs in the zebrafish larvae showed no enriched expression of *ahr1a* or *ahr1b* (**Fig. S1C, D**). However, we found that EECs did express high levels of the sensory channel *trpa1b* (**Fig. S1E**), which can be activated by tryptophan metabolites (*19*). To determine whether Trpa1 activation induces *il22* expression in EECs, we micro-injected tryptophan metabolite indole into the guts of *il22*:mCherry; *neurod1*:GFP double transgenic larvae (**Fig. 1E**). Live imaging showed an increase of *il22*-expressing EECs upon indole injection (**Fig. 1F, G**). Furthermore, treating larvae with a Trpa1 receptor inhibitor reduced *il22* expression in larval guts (**Fig. 1H, I**). Together, our data reveal that activation of Trpa1, likely stimulated via microbial metabolites, promotes *il22* expression in larval gut EECs.

### Zebrafish IL-22 promotes gut immunity and shapes gut microbiota composition

Having found that IL-22 is primarily expressed in larval EECs rather than in immune cells, we then investigated its function at this stage of development. To infer IL-22-regulated biological processes, we used CRISPR/Cas9 to generate *il22*^-/-^ mutant zebrafish (**Fig. S2A-D**). The *il22* mutant fish did not show gross morphological defects, reached adulthood, and were fertile. (**Fig. S2E-I**). We took an unbiased approach and performed bulk RNA-sequencing of guts from WT and *il22*^-/-^ larvae (**Fig. 2A-C and S2J, K**). 917 transcripts were significantly down-regulated while 1050 were up-regulated in *il22*^-/-^ larval guts (**Fig. 2B, Table S3**). Gene ontology (GO) analysis indicated that most down-regulated genes in *il22*^-/-^ were associated with response to bacteria (**Fig. 2C**). In mice, IL-22 released from lymphocytes promotes anti-bacterial gene expression in the gut epithelium, shaping microbiota composition and protecting from bacterial infection (*13, 23, 25*). To investigate whether IL-22 influences microbiota composition in the developing zebrafish, we performed 16S RNA sequencing on guts from WT and *il22*^-/-^ larvae (**Fig. 2D-G, Table S4**). RDA analysis revealed consistent clustering of biological replicates, with a distinct separation between WT and *il22*^-/-^ larvae, pointing to reproducible differences in their bacterial communities (**Fig. 2D**). The Shannon diversity index indicated a comparable level of microbial diversity between the two groups (**Fig. 2E)**. Significant differences were observed at the family level, with an enrichment of *Staphylococcaceae* and *Enterobacteriaceae*, as well as a reduction of *Lactobacilleae* in *il22*^-/-^ larvae (**Fig. 2F, G**). These findings parallel the function of IL-22 in repressing *Enterobacteriaceae* and promoting *Lactobacilleae* in mice through antimicrobial peptide induction (*26*). Finally, to confirm the gut protective role of zebrafish *il22*, we performed bath infection with live *Edwardsiella tarda*, a well-known fish gut pathogen (**Fig. S3A, B**). *il22*^-/-^ showed an increased susceptibility to infection with this bacterium. Collectively, our results identify a conserved role for IL-22 in modulating microbiota composition and safeguarding against gut bacterial infection, despite being primarily produced by epithelial cells instead of immune cells in the developing gut.

**Figure 2.**
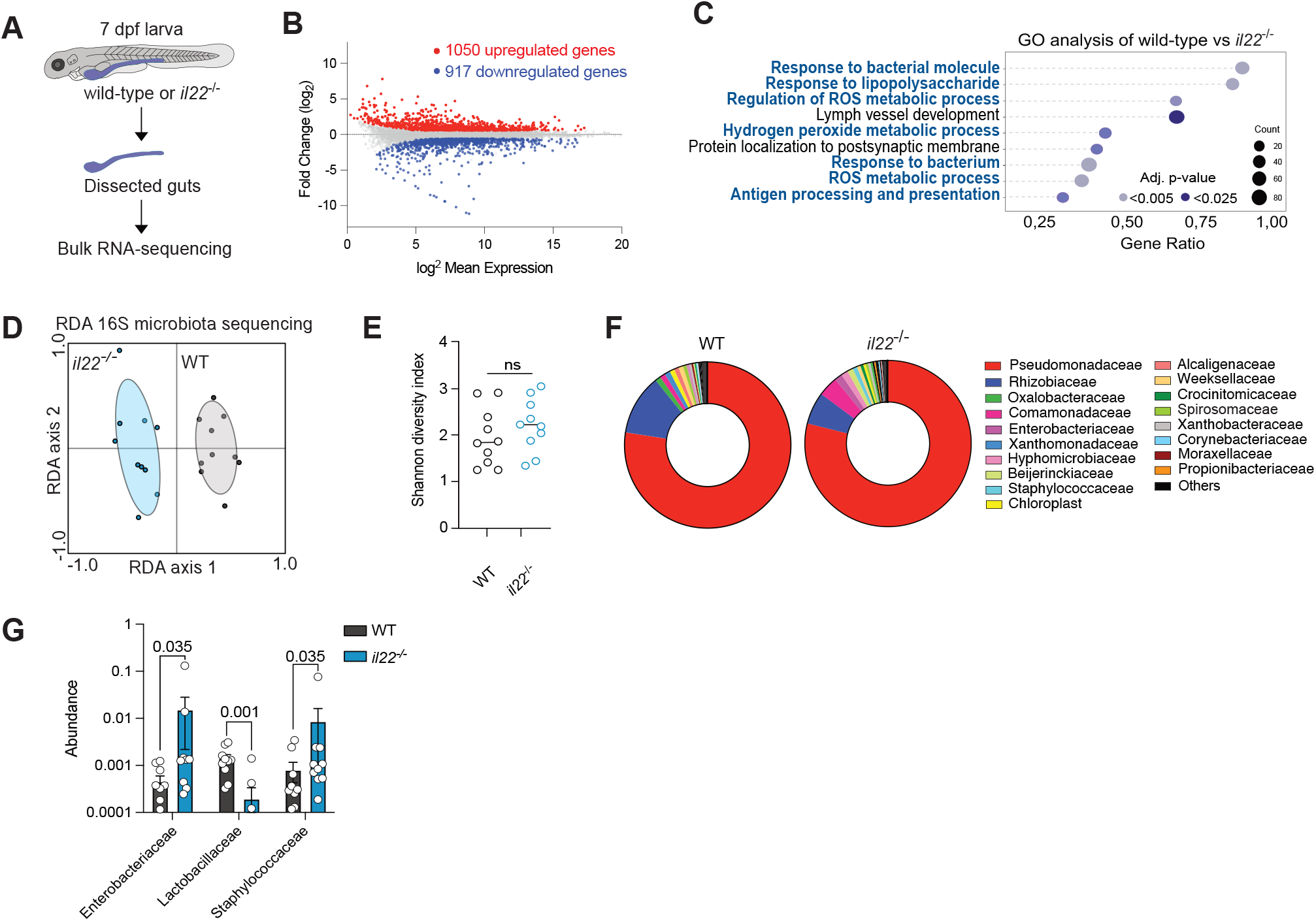
Zebrafish IL-22 shapes gut microbiota composition in larvae. **A**. Diagram showing gut dissection of WT or *il22*^-/-^ larvae at 7 dpf for bulk RNA-sequencing. **B**. MA plot showing the dysregulated genes in guts of *il22*^-/-^ compared to WT larvae with a *P* < 0,05 and a fold change > 1.5. The 917 downregulated genes are labeled in blue and the 1050 upregulated genes are shown in red. **C**. Gene ontology analysis of genes downregulated in *il22*^-/-^. Most terms are associated with anti-bacterial immunity. **D**. RDA plot of 16S rRNA-sequencing comparing 7 dpf WT and *il22*^-/-^ larval intestines. **E**. Shannon-Wiener microbiota diversity using 16S rRNA sequencing between WT and *il22*^-/-^ guts. **F**. Taxon-based analysis at the family level between WT and *il22*^-/-^. **G**. Abundance of the indicated bacterial families in guts of WT and *il22*^-/-^ larvae. Statistical analysis was performed with multiple comparisons 2-way ANOVA ** *P* < 0,01, *** *P* < 0,001. Data are representative of at least two independent experiments (except RNA-sequencing for which we had 4 replicates of each condition).

### Zebrafish IL-22 modulates enteric nervous system function and gut motility

We observed the upregulation of 1050 genes in *il22*^-/-^ compared to WT guts (**Fig. 2B, Table S3**). Unexpectedly, GO analysis showed enrichment of genes associated with neuronal and axon development (**Fig. 3A, B**). Therefore, we sought to understand how IL-22 influences the enteric nervous system. Overall neuron numbers did not appear to be affected (**Fig. 3C, D**), however, dysregulation of genes linked to ion transport and calcium signaling (**Fig. 3B**) suggested impaired neuronal activity in *il22*^*-/-*^ larvae. To assess changes in activity, we performed calcium imaging of enteric neurons in live zebrafish larvae using a neuron-specific GCaMP6f reporter (HuC:GCaMP6f) and light-sheet microscopy (**Fig. 3E, Movie S1, 2**) (*27*). We observed altered gut neuronal activity in *il22*^-/-^ larvae, which was characterized by changes in the periodicity of calcium activity (**Fig. 3F-H**). As a key function of enteric neurons is to regulate gut motility, we looked at whether the reduced neuronal activity could be linked to reduced gut motility. Live imaging revealed both reduced gut motility (**Fig. 3I, J, Movie S3-6**) and slower food transit in *il22*^-/-^ larvae (**Fig. 3K, L**). Together, our results reveal a role of EEC-derived IL-22 in the regulation of enteric nervous system function and gut motility.

**Figure 3.**
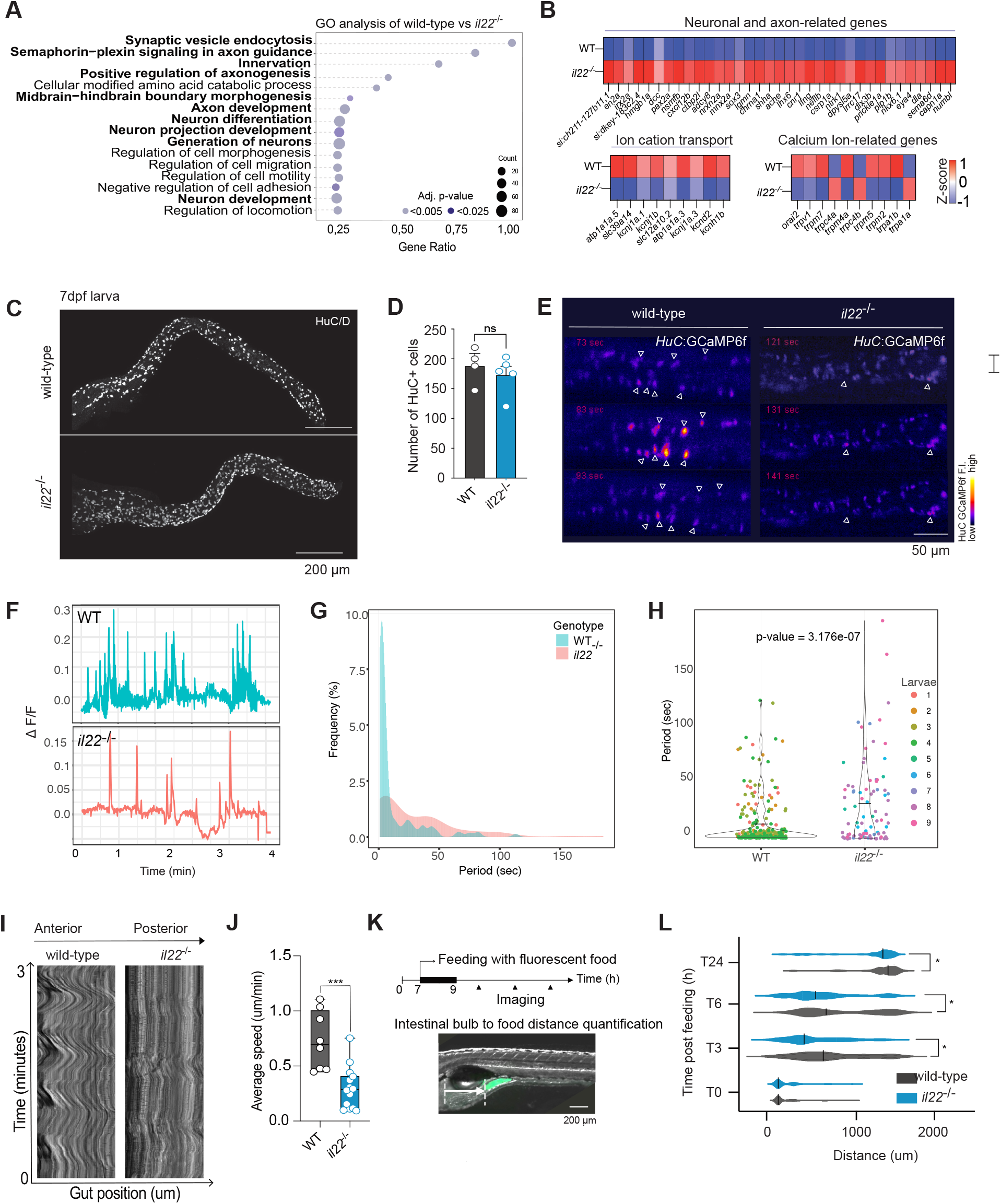
IL-22 controls enteric neuron activity and gut motility. **A**. Gene ontology analysis comparing WT and *il22*^-/-^ upregulated genes. GO terms in bold are associated with neuronal development. **B**. Heatmaps comparing gene expression levels in guts of WT and *il22*. **C**. Image of WT and *il22*^-/-^ guts of larvae stained with an anti-HuC/D (a pan-neuronal marker) antibody. Scale bar = 200um. **D**. Quantification of the number of HuC/D positive cells in the gut of 7 dpf WT and *il22*^-/-^. **E**. Snapshots of enteric neuron activity imaging using light-sheet microscopy of the Tg(*Huc*:Gcamp6f) transgenic line in 7 dpf WT and *il22*^-/-^ larvae. Triangles indicate segmented neurons (see methods). **F**. Examples of neuronal activity temporal traces showing the change in fluorescence intensity relative to the baseline (ΔF/F) in guts of 7 dpf WT and *il22*^-/-^ larvae. **G**. Histogram of the time distribution between consecutive peaks of neuron activation (period) in guts of 7 dpf WT and *il22*^-/-^ larvae. **H**. Comparison of period composition of gut neural trace activation between 7 dpf WT and *il22*^-/-^ larvae. Wilcoxon rank-sum test (p-value = 3.17e-07). **I**. Kymograph analysis 180 um in the midgut of 7 dpf WT and *il22*^-/-^ larvae using ImageJ. **J**. Gut motility speed of 7 dpf WT and *il22*^-/-^ larvae. **K**. Schematic of food transit experiment. In brief, larvae were fed for 2 hours with dry food coupled with fluorescent beads, and the distance from the anterior bulb to the fluorescent food was measured at different time points. **L**. Location of the fluorescent food in the intestine of WT and *il22*^-/-^ larvae at different time points. Statistical analysis was performed with unpaired T-test, ns: not significant, * *P* < 0,05, ** *P* < 0,01, *** *P* < 0,001. Data are representative of at least two independent experiments (except RNA-sequencing for which we had 4 replicates of each condition).

### IL-22 signaling in gut epithelial cells is sufficient to sustain gut motility

Gut motility is influenced by the central nervous system and other organs. Hence, we aimed to determine if IL-22 signaling in the gut can sustain gut motility. To this end, we sought to create a zebrafish line expressing the IL-22 receptor exclusively in gut epithelial cells. Based on synteny, we identified the zebrafish *crfb14* (cytokine receptor family member B14) as a putative ortholog of the human IL-22 receptor-specific gene *IL22RA1. crfb14* showed higher expression in gut epithelial cells than non-epithelial cells (**Fig. S4A, B**). We generated a *crfb14* mutant line that failed to induce IL-22 target gene expression and had impaired gut motility, similar to *il22*^-/-^ larvae (**Fig. S4C-G, Movie 7**). Analysis of public scRNA-seq datasets of zebrafish larval intestines (*21*) revealed *crfb14* expression mainly in gut epithelial cells and not in enteric neurons or smooth muscles, both key cell types controlling gut motility (**Fig. S4H**). As Crfb14 was essential for gut motility, we next asked whether restoring its expression solely in epithelial cells could correct the motility defects. We generated a transgenic line expressing *crfb14* exclusively in IECs by driving *crfb14* expression with the gut epithelial-specific *cldn15la* promoter in *crfb14*^*-/-*^ fish (*crfb14*^*-/-*^ ;*cldn15la:crfb14*, from now on IEC-*crfb14* **Fig. S4I-K**). Expressing *crfb14* in IECs alone restored gut motility (**Fig. S4L, M**), indicating that epithelial IL-22 signaling is sufficient to sustain gut motility. Altogether, we identified the gene encoding the IL-22-specific receptor chain in zebrafish and demonstrated that IL-22 controls gut motility through gut epithelial cells.

### Intestinal motility impairment in *il22*^*-/-*^ larvae is restored by co-housing and live microbiota transfer from WT larvae

We observed an altered microbiota composition in *il22*^-/-^ larvae. Since the gut microbiota is crucial for the development and function of enteric neurons that drive gut motility (*28*), we asked whether the IL-22 influence on gut motility could be related to these alterations. We first analyzed the gut motility of *il22*^-/-^ germ-free (GF) larvae to their conventionally reared (CV) counterparts. The gut motility impairment in *il22*^-/-^ was consistent across GF and CV conditions (**Fig. 4B, S5A**). To determine whether the microbiota could play a role in the observed defects, we co-housed *il22*^-/-^ GF with WT CV larvae to promote gut microbiota transfer from WT to mutants (*13, 29*) (**Fig. 4A, B, S5A**). Interestingly, the impaired gut motility of *il22*^-/-^ was restored when co-housed with WT larvae. These results suggested that factors from WT larvae can compensate for the lack of IL-22. To determine whether live bacteria or molecules present in the medium of WT larvae were responsible for this compensation, we transferred the medium from WT CV to *il22*^-/-^ GF larvae (**Fig. 4C**). Interestingly, this transfer restored the gut motility impairment of *il22*^-/-^. However, this did not occur if the medium from WT larvae was pretreated with antibiotics, indicating a critical role of live bacteria in the rescue process (**Fig. 4C, D, S5B**). Collectively, our data show that live bacteria from WT larvae can efficiently rescue the gut motility defect of *il22*^-/-^ larvae, indicating that the role of IL-22 in early-life gut physiology involves regulating the microbiota.

**Figure 4.**
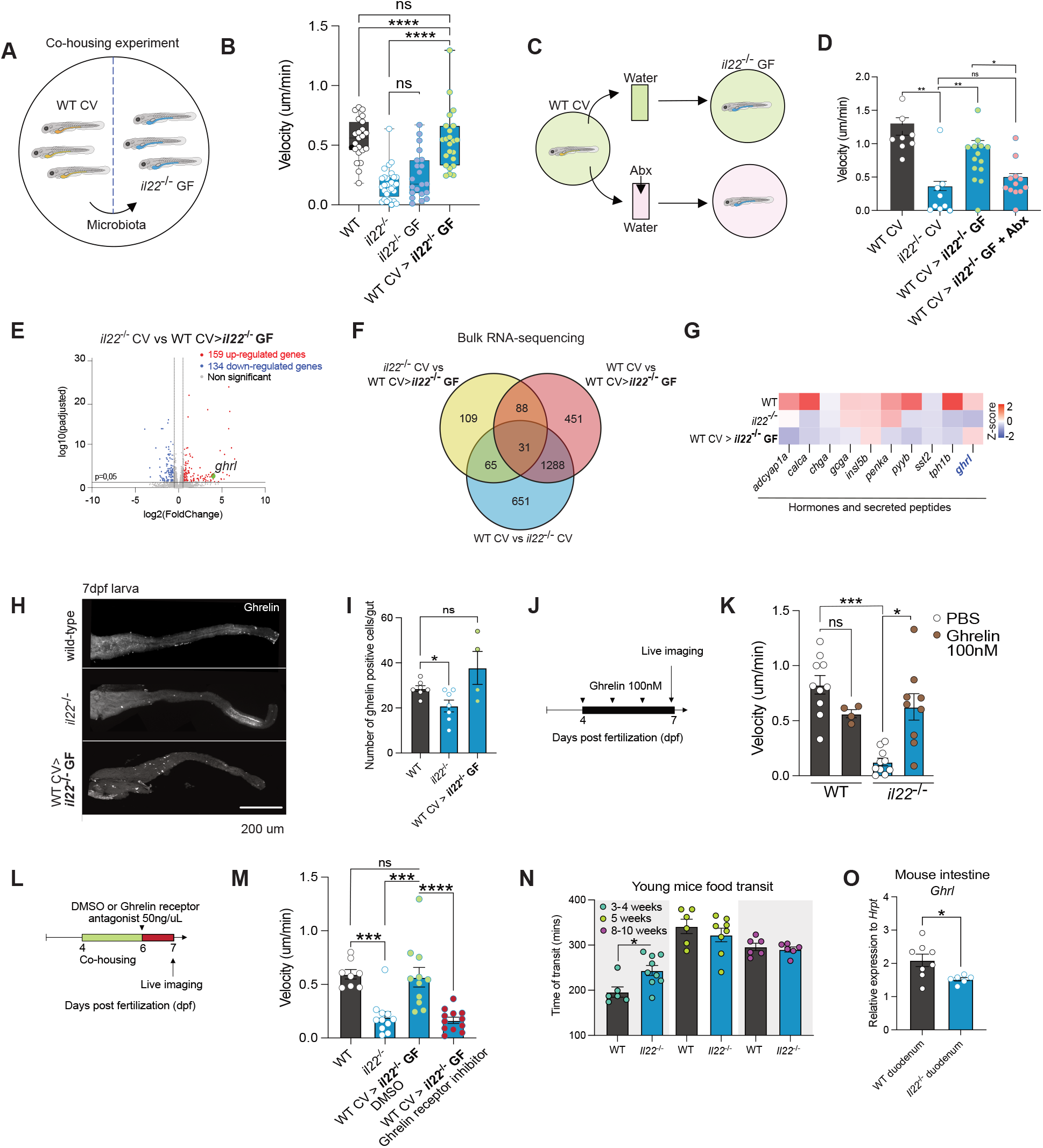
Ghrelin mediates IL-22-dependent gut motility. **A**. Schematic of the co-housing protocol. Briefly, conventionally-reared (CV) larvae were co-housed in the same petri dish with GF larvae for three consecutive days to promote microbiota transfer followed by gut motility analysis. **B**. Gut motility measurements of larvae of the indicated genotypes in CV and GF conditions, as well as co-housing. The genotypes of the larvae in which gut motility was measured are shown in bold. **C**. Schematic of medium transfer experiment. Fresh medium from WT CV was transferred to *il22*^-/-^ GF larvae every day for 3 consecutive days without or with antibiotics (Abx) treatment. **D**. Gut motility speed in larvae of the indicated genotypes in CV conditions as well as in *il22*^-/-^ GF larvae receiving medium from WT CV or WT CV treated with antibiotics. **E**. Volcano plot showing downregulated genes (blue), upregulated genes (red), and unchanged genes (gray) in guts of *il22*^-/-^ CV compared to *il22*^-/-^ GF co-housed with WT CV larvae. **F**. Triple Venn diagram displays the number of differentially expressed genes across the indicated comparisons. **G**. Heatmap comparing levels of expression of the indicated genes in guts of WT CV, *il22*^-/-^ CV, and *il22*^-/-^ GF co-housed with WT CV larvae. **H**. Representative pictures of immunofluorescence on dissected guts of WT CV, *il22*^-/-^ CV, and *il22*^-/-^ GF co-housed with WT CV larvae. **I**. Quantification of numbers of Ghrelin-positive cells after immunofluorescence on dissected guts of WT CV, *il22*^-/-^ CV, and *il22*^-/-^ GF co-housed with WT CV larvae. **J**. Schematic describing the treatment of larvae with Ghrelin. Ghrelin protein (100nM) was added every day from 4 to 7 dpf to the medium of WT or *il22*^-/-^ larvae. Gut motility was analyzed by live imaging at 7 dpf. **K**. Gut motility speed quantification of WT, and *il22*^-/-^ treated with PBS or Ghrelin. **L**. Schematic describing the treatment of larvae with a Ghrelin receptor antagonist (50 ng/uL) during the last 4 hours of a 3-day co-housing period between *il22*^-/-^ GF with WT CV larvae. **M**. Gut motility speed quantification of WT CV, *il22*^-/-^ CV, *il22*^-/-^ GF co-housed with WT CV larvae treated with DMSO, and *il22*^-/-^ GF co-housed with WT CV larvae treated with a Ghrelin receptor inhibitor. **N**. Food transit experiment in 3-4, 5, and 8-10 weeks old WT or *Il22*^-/-^ mice. **O**. RT-qPCR on the duodenum of 3 weeks old WT and *Il22*^-/-^ mice. Statistical analysis was performed with Mann-Whitney: * *P* < 0,05, ** *P* < 0,01, *** *P* < 0,001, **** *P* < 0,0001. These data are representative of at least two independent experiments.

### Microbiota transfer rescues *il22*^*-/-*^ motility impairment via ghrelin

To better understand what factors are involved in the gut motility rescue, we investigated the transcriptional changes in *il22*^-/-^ guts upon co-housing with WT larvae. We performed bulk RNA-sequencing on dissected guts from WT CV and *il22*^-/-^ CV larvae, as well as *il22*^-/-^ GF co-housed with WT CV larvae (**Fig. S5C, D, Table S5-8**). WT CV compared to *il22*^-/-^ co-housed with WT larvae showed a similar number of differentially expressed genes than WT CV compared to *il22*^-/-^ CV, still with enrichment of GO terms related to immune responses and neuronal development (**Fig. S5E, F, Table S5**). These results suggest that the expression of these genes is not significantly impacted in *il22*^-/-^ upon microbiota transfer from WT larvae. Indeed, comparing *il22*^-/-^ CV with *il22*^-/-^ co-housed with WT larvae revealed fewer than 300 genes with different expression levels (**Fig. 4E)**. Among these, we observed upregulation of *ghrl*, which encodes ghrelin, a hormone that regulates satiety, food intake, and intestinal motility (*30, 31*) (**Fig. 4E, Table S7**). Furthermore, from the genes dysregulated in *il22*^-/-^ CV guts compared to WT CV, co-housing the mutants with WT restored the expression of only 65 genes to WT CV levels, including *ghrl* (**Fig. 4F, Table S8**).

Additionally, we found that compared to WT, *il22*^-/-^ guts had dysregulated expression of several EEC genes, including hormones and neuropeptides involved in gut motility control (**Fig. 4G**). From these genes, only *ghrl* showed a significant rescue of expression levels when co-housing *il22*^-/-^ with WT larvae, which was confirmed by immunostaining in dissected guts (**Fig. 4G-I**). We thus wondered whether ghrelin played a role in mediating gut motility rescue in *il22*^-/-^ larvae upon co-housing with WT. Treating *il22*^-/-^ larvae with ghrelin peptide was able to restore intestinal motility defects (**Fig. 4J, K, S5G**). Moreover, treatment with a ghrelin receptor inhibitor impeded the gut motility rescue of *il22*^-/-^ GF larvae when co-housed with WT CV (**Fig. 4L, M, S5H)**. These findings indicate that ghrelin plays an important role in regulating intestinal motility in *il22*^-/-^ and mediates the improved gut motility upon co-housing with WT larvae.

### IL-22 promotes gut motility in mice during early life

To address whether our findings in zebrafish larvae can be translated to mammals, we measured food transit speed in WT or *Il22*^-/-^ mice at three different stages (3-4, 5, or 8-10 weeks old). Interestingly, we observed a slower food transit time in *Il22*^-/-^ at 3-4 weeks (**Fig. 4N**). Notably, this impairment did not take place at older ages, indicating that IL-22 influences gut motility in mice only during early life. To determine whether *Il22*^-/-^ mice exhibit a comparable deficiency in hormone expression, we performed qPCR on the intestines of 3-week-old mice and observed reduced expression levels of *Ghrelin* in *Il22*^-/-^ mice relative to WT (**Fig. 4O**). These data reveal an evolutionarily conserved role for IL-22 in modulating gut function during the early life of vertebrates.

## Discussion

Our study identifies a circuit in which IL-22-microbiota interactions via EECs control intestinal physiology during development (**Figure S6**). We demonstrated that IL-22 is expressed and functional in zebrafish larval EECs before lymphocytes, such as ILCs and T cells—known producers of this cytokine in adult zebrafish and mammals—populate the gut. Recent studies have shown that during early vertebrate development, epithelial cells at the blastula stage of zebrafish and mice can perform phagocytic immune functions before immune cells emerge (*32*). Together with our findings, these data support the concept that epithelial cells in developing organisms can perform functions traditionally attributed to immune cells.

Our data suggest a similar role of microbiota-derived metabolites in inducing IL-22 production in zebrafish and mammals. In mice, tryptophan regulates gut homeostasis through IL-22 produced by lymphocytes upon AhR activation (*33*). Bacterial-derived tryptophan also activates EECs in vertebrates and invertebrates (*34, 35*), inducing hormone expression and gut motility through Trpa1 activation in zebrafish (*19*). We show that IL-22 expression is regulated by tryptophan metabolites in EECs, suggesting an evolutionarily conserved pathway involving the Trpa1 receptor in zebrafish epithelial cells and AhR in mammalian lymphocytes. Trpa1 senses pain (*36*), heat (*37*) and mechanical stress (*36, 38*); thus, our work suggests additional functions of IL-22 upon these stimuli.

The imbalance of gut microbiota in *il22*^-/-^ zebrafish parallels observations in *Il22*^-/-^ mice (*13*), highlighting the conserved role of this cytokine in microbiota composition. In mice, we found a role of IL-22 on gut motility during the stage coinciding with weaning, which is marked by significant shifts in both the microbiota and the immune system (*4*). Additionally, we revealed a new role of IL-22 in the function of the enteric nervous system and gut motility in the developing zebrafish. Together, our data suggest an evolutionary conserved role of IL-22 in gut motility in early life.

We demonstrate a key role for microbiota transfer in restoring intestinal physiology in IL-22-deficient zebrafish. Furthermore, we identified the peptide hormone ghrelin as the factor mediating the rescue of gut motility impairment in *il22*^-/-^ zebrafish upon microbiota transfer from WT larvae. Ghrelin is known to stimulate motility and gastric emptying *via* the vagus nerve, relaying nutritional information along the gut-brain axis (*31*). The role of this axis in the observed rescue as well as the systemic effects of impaired gut hormone expression in IL-22-deficient animals remain to be addressed. Ghrelin expression is regulated by the microbiota (*39*), which aligns with our findings and suggests a circuit where IL-22 controls microbiota composition to ensure proper ghrelin expression and gut function during vertebrate development.

Collectively, our findings reveal key mechanisms by which host-microbiota interactions during development, mediated by a cytokine in epithelial cells, influence intestinal physiology before a mature immune system is established. This advances our understanding of early life gut physiology, and also sets the basis for exploring these mechanisms in other developing tissues, as well as in inter-organ and inter-individual communication.

## Acknowledgments

The authors would like to acknowledge the Cell and Tissue Imaging Platform – PICT-IBiSA (member of France–Bioimaging – ANR-10-INBS-04) of the U934/UMR3215 Unit of the Institut Curie for help with microscopy, in particular to Olivier Leroy for his help for analyzing Calcium activity. We also thank the members of the animal facility of the Institut Curie for zebrafish care. Life Science Editors provided editing services for this manuscript.

## Funding

This work was supported by the Institut Curie, INSERM, and CNRS, and the grants listed below.

Ville de Paris emergence program (2020 DAE 78) (PPH)

FRM amorçage (AJE201905008718) (PPH, AM)

ATIP-Avenir starting grant R21045DS (PPH, AM)

ERC-StG Cytok-Gut 101041422 (PPH, SR, AJK, LM)

ANR IPOGUT (ANR-22-CE17-0040) and INCEPTION (ANR-20-CE16-0020) (FDB)

PhD fellowships from La ligue contre le cancer (IP/SC-16060) and la Fondation pour la recherche médicale (FDT202204014884) supported SR.

The PhD fellowships from the Ministère de l’Enseignement Supérieur et de la Recherche and from the Fondation pour la Recherche Médicale (FDT202304016654) supported YS.

IMY., has received funding from the European Union’s Horizon 2020 research and innovation programme under the Marie Skłodowska-Curie grant agreement No 847718 (Institut Curie EuReCa PhD Programme). This publication reflects only the author’s view and that the European Research Agency is not responsible for any use that may be made of the information it contains.

Laboratoire d’Excellence (Labex) DEEP (ANR-11-LBX-0044, ANR-10-IDEX-0001-02 PSL).

## Author contributions

P.P.H., and S.R. conceived the project and designed the experiments with inputs from Y.S., I.MY., A.M., G.G., C.BG., P.D., A.JK. F.G., and G.E. performed the mouse experiments at Institut Pasteur, Paris. E.M., and G.S., performed Calcium imaging in the Ecole Normale Supérieure, Paris. L.M., performed qPCR on bacterial samples. A.JK. analyzed the scRNAseq dataset from *Nayar* et al, 2021. S.B., and J.B. performed and analyzed the 16S rRNA-sequencing experiments at Wageningen University, Netherlands. V.DD., and F.DB. generated the *crfb14* zebrafish mutant at the Institut Curie, Paris. JP.L., G.L., and P.P.H. generated the *il22* mutant and *il22*:mCherry zebrafish lines. K.QC., and C.F. generated the IEC-*crfb14* line, performed live imaging, and conducted bacterial infection and quantified larval survival at Andres Bello University in Chile. P.P.H., and S.R. wrote the manuscript with assistance from the remaining authors.

## Data and materials availability

The new reagents and datasets generated during the current study are available from the corresponding author upon request.

