## Supplementary materials for "Interleukin-22 in enteroendocrine cells controls early-life gut motility through interactions with the microbiota"

### The PDF file includes:

Materials and Methods
Figs. S1 to S6
Tables S1
References

### Materials and Methods

**Zebrafish lines and husbandry** The maintenance of zebrafish wild-type line (AB), transgenic lines Tg(*mpx*:EGFP)*114* (*40*), Tg(*neurod1*:GFP) (*22*), TgBAC(*cldn15la*:GFP)*pd1034* (*20*), Tg(*neurod1*:GCaMP6f)*icm05* (*27*) and the newly generated lines was performed in accordance with European Union regulations on laboratory animals using protocol numbers : APAFIS#27495-2020100614519712 v14.2.2, #2019_010 and #2022-008 (approved the French Ministry of Research). Zebrafish larvae used in foreign collaborations abroad were maintained according to the "Acta de Aprobación 004/2021" provided by the Universidad Andrés Bello, Chile.

**Mice** Wild-type (C57BL/6) and *Il22*$^{-/-}$ mice were maintained in the Institut Pasteur animal facilities. Animal care and experiments were performed according the committee on animal experimentation of the Institut Pasteur and authorized by the French Ministry of Research.

**Construction of mutant zebrafish lines and genotyping** The zebrafish coding sequence for the ortholog of the human *IL22* gene (Gene name: *il22*, ENSEMBL ID: ENSDARG00000045673) was targeted by CRISPR/Cas9 with specific sgRNA: CTGTGCTCGTGCTTTTTGAG. The same method was used for the *crfb14* gene in zebrafish that we identified as the IL22RA1 ortholog (Gene name: *ifnlr1*, ENSEMBL ID: ENSDARG00000087131) with this specific sgRNA: TCAAACGGCTCTTT. For the maintenance of WT, *il22*$^{-/-}$ and *crfb14*$^{-/-}$ mutant stocks, zebrafish embryos derived from the incrossing of heterozygous *il22* and *crfb14* individuals were reared and genotyped when they reached adulthood by cutting off part of the caudal fin (Fin Clip). Briefly, Adult and larvae zebrafish were anesthetized with tricaine (100 ug/ml, Sigma Cat#A5040), their tails were cut and incubated during 1h at 56°C with FinClip buffer (Tris pH 8 10mM, EDTA 10mM, NaCl 200mM, SDS 0,5%) containing Proteinase K (0,2mg/mL, Invitrogen #25530-049). The mutation was genotyped using the KASP system (LGC genomics). The KASP assay was performed according to the manufacturer's instructions (*41*). The reaction mix per reaction consists of 5.1 µl KASP Master Mix containing the two allele-specific primers and one reverse primer (see Table 1) and 0.138 µl Assay Mix containing universal fluorescent probes, Taq polymerase and dNTPs in an optimized buffer solution. The only exception was for *crfb14*$^{-/-}$ that we genotyped by RT-qPCR (see primers Table 1).

**Construction of the *il22:mCherry* transgenic zebrafish line** The 6.5 kb SpeI-PstI fragment from PAC clone BUSMP706A0151Q01 (IMAGENE) covering the *il22* promoter was cloned ahead of the ORF for a farnesylated version of mCherry in a Tol2 derivative vector to yield vector pTol2-p*il22*mC-F. The fragment includes exon 1 including the first codons of the zebrafish *il22* ORF. This construct was co-injected with Tol2 mRNA into 1-cell stage eggs of AB origin. Screening for mCherry positive fish was performed by PCR.

**Construction of the *cldn15la:GFP-p2a-crfb14* zebrafish line** The original plasmid was the pDestTol2pA2_349cldn15la-GFP-KRASV12 kindly provided by Filippo Del Bene. The *crfb14* coding sequence (Gene name: *ifnlr1*, ENSEMBL ID: ENSDARG00000087131) was synthesized by Genescript and replaced the *kras*$^{v12}$ gene expression in the aforementioned plasmid. The generated plasmid construct (25 ng/µl) was then co-injected with mRNA transposase (50 ng/µl) into 1-cell stage *crfb14*$^{-/-}$ embryos (*42*) and the resulting embryos were grown to adulthood for stable line screening. The rescue line name has been shortened to *crfb14*$^{-/-}$ F1.

**Length and area measurements** Larvae were anesthetized with tricaine and mounted in 3% methylcellulose for live imaging. Body and intestinal length measurements were performed from the intestinal bulb until the end of the intestine at the anal pore of the larva, and body length or area measurements in adults and larvae were done from the head to the tail. The measurements were quantified using ImageJ software (NIH).

**Alcian blue staining** Fixed larvae in Paraformaldehyde 4% (Polysciences inc., #04018-1) were rinsed with acidic ethanol (70% ethanol (VWR, #20821.310) with 1% concentrated hydrocholoric acid (AnalaR NORMARK, #20252.290)) before being incubated with 0.1% alcian blue (Sigma, #33864-99-2) diluted in 80% ethanol, 20% glacial acetic acid (SAFC, #ARK2183) during 3h at RT. Then, larvae were washed with acidic ethanol and imaged with an Upright Epifluorescence Microscope (Leica DM4 B) equipped with a color camera (DFC4500 Leica).

**Immunofluorescence staining and imaging** *Whole larvae* : Immunostaining was performed on whole larvae at 5 or 7 days post fertilization. Paraformaldehyde at 4% was used to fix zebrafish larvae overnight at 4°C. The sample were then washed with distilled water. Fixed larvae were then permeabilized with cold 100% acetone (Honeywell, #32213) during 20 min at 4°C before being washed three times with PBST (PBS 1X + 0,5% Triton-X100 (Invitrogen, #10717503). Samples were then permeabilized with 1mg/mL Collagenase from *Clostridium histolyticum* (Sigma, #C2139) during 2h at room temperature. Samples were then washed with PBST and blocked with 10% of FBS (fetal bovine serum)/PBST at room temperature for more than 2h. The primary antibodies (see Table 1) were diluted in blocking solution solution and incubated at 4°C for more than 24h. Following primary antibody incubation, the samples were washed with PBST solution and incubated during at least 2h with secondary antibodies (see Table 1) at room temperature in the dark.
*Dissected guts* : Immunostaining was performed on dissected guts at 7 days post fertilization. Paraformaldehyde at 4% was used to fix zebrafish larvae dissected intestines 1h at room temperature. The samples were then washed with distilled water and incubated overnight with the primary antibodies (see Table 1) diluted in PBS 1X. Then, samples were washed with PBS 1X + 0,5% Triton-X100 (Invitrogen, #10717503) three times before being incubated during at least 2h with secondary antibodies (see Table 1) at room temperature in the dark. Dissected guts were then washed in PBS twice and mounted with mounting medium containing DAPI (Thermofisher Scientific, #P36931).
Imaging was performed with THUNDER Imager Model Organism (Leica) with lens and analyzed with ImageJ software.

**Neuronal activity measurements** To characterize the neuronal activity of Tg(*HuC*:GCaMP6f), larvae were first embedded laterally in a thin layer of 4% Low melting point agarose (Promega, #V2111). Then, we used selective-plane illumination microscopy (SPIM) to record the neuronal activity at cellular resolution across the gut. Optical sectioning was achieved by the generation of a micrometer-thick light sheet to excite GCaMP from the side and the front of the larva. The GCaMP emission was collected by a camera whose optical axis was orthogonal to the excitation plane (a 488 nm laser, Phoxx 480-200, Omicron). In both arms the laser beam was first filtered by a 488 cleanup filter (F488 Omicron) and coupled to a single-mode fiber optic. The beam was expanded using a telescope (f = 50 mm, LA1131-A, and f = 150 mm, LA1433-A, Thorlabs) and projected onto two orthogonal galvanometric mirrors (HP 6215H Cambridge technology) to scan the laser beam, whose angular displacement were converted into position displacement by a scan lens (f = 75 mm AC508-075-A-ML, Thorlabs). The laser beams were then refocused by a tube lens (f = 180 mm, U-TLUIR, Olympus) and focused on the pupil of a low-NA (0.16) 5x objective lens (UPlan SAPO 4x, NA = 0.16, Olympus) facing the specimen chamber. The arrangement yielded a 1mm-wide illumination sheet and a beam waist of 3.2mm (1/e2). The emitted fluorescence light was collected by a high-NA water-dipping objective (N16XLWD-PF, 16x, NA = 0.8, Nikon) mounted vertically on a piezo translation stage (PI PZ222E). A tube lens (f = 180mm U-TR30IR, Olympus), a notch filter (NF03-488, to filter the laser's excitation light), a band-pass filter (FF01 525/50 Semrock) and a low-pass filter (FF01 680 SP25 Semrock, to filter the IR light) were used to create an image of the GCaMP emitted fluorescence on a sCMOS sensor (Orca Flash 4.0, Hamamatsu). The volumetric gut recordings were obtained by sequentially recording the fluorescence in 40 coronal sections spaced by 3 um. For this purpose, the light sheet was scanned vertically in the dorso-ventral direction in synchrony with the objective of the emission path. The camera was triggered to acquire an image every Texposure = 10 ms. Once the 40 coronal sections were recorded, the position of the light sheet and the objective of the emission path was reset to their initial dorsal position (Treset = 100 ms). This resulted in a volumetric acquisition time of 0.5 s or a rate of 2 Hz. The cell tracking and intensity measurement were performed using Imaris.

**Body-intestine dissection in zebrafish larva** Larvae at 7dpf were euthanized by overdosing them with tricaine. Intestines were extracted mechanically by using tweezers (WPI, #142400). 5 intestines and their respective bodies were used for each replicates and further analyzed by RT-qPCR.

**Quantitative real-time PCR** *Zebrafish* : Intestines or body carcasses were pooled and RNA was extracted using the Single cell RNA purification kit (Norgen Biotek Corp, Cat. 51800) following manufacturer's instructions. Synthesis of cDNA was performed using the M-MLV Reverse Transcriptase Kit (Invitrogen). Real-time PCR was performed using the Rox SYBR Green MasterMix dTTP Blue Kit (Takyon) and run on a Thermo ABI ViiA 7 Real-Time PCR System (Thermo Applied Biosystems).
*Mice* : For epithelial cell isolation, the small intestine was collected, and the tissue was cleaned from feces and fat. The epithelial layer was dissociated by incubating the cleaned tissue for 15 min in HBSS supplemented with 5 mM EDTA and 10 mM HEPES at 37°C (twice). 1 mL of the epithelial cell suspension was collected after each round of dissociation. The cells were pelleted and resuspended in 1ml of Trizol (Life Technologies). The cells were homogenized using Precellys Evolution homogenizer (Bertin technologies).The RNA from epithelium and was extracted according to the PureLink RNAminikit (Thermo Fisher) protocol and cDNA was transcribed using the Superscript IV Reverse Transcription Kit (Thermofisher). Real-time PCR was performed using the Rox SYBR Green MasterMix dTTP Blue Kit (Takyon) and run on a Thermo ABI ViiA 7 Real-Time PCR System (Thermo Applied Biosystems).

Samples were analyzed using ΔCt method. The mean Ct value of housekeeping gene (*ef1a*) was used for normalization. Primers used for qRT-PCR are found in Table 1.

**Fluorescence-activated cell sorting (FACS)** To acquire the intestinal epithelial population, approximately 100 7 days post-fertilization TgBAC(*cldn15la*:GFP) zebrafish larvae were collected, then intestines were dissected and placed into PBS on ice with a dissection time of maximum 2 hours. Intestinal cell dissociation was performed using TrypLE Express (Gibco, #12605028) for 1h at 37°C, pipetting up and down every 10 minutes to support digestion. Digested samples were spun at 1500g for 5 min at 4°C and wash twice with PBS 1X before being resuspended together with PBS 1X and 10% FBS (fetal bovine serum). Filtered cells were immediately subjected to FACS at the Institut Curie Flow Cytometry Platform with a Sony SH800 Cell Sorter. Dead cells were excluded from analysis using a combination of Calcein Blue (Invitrogen, #65-0855-39) and Propidium Iodide viability stains (Sigma, #P4864). Non-transgenic and single transgenic controls (pools of 10 dissected guts) were prepared as above and used for gating and compensation. RNA isolation was done using on average 30000 GFP+ or GFP- sorted cells with the Single Cell RNA Purification Kit from Norgen Biotek Corp and reverse transcribed using Superscript IV Reverse Transcriptase (Life Technologies, #18090050) following manufacturer's instructions. Quantitative PCR was performed with gene-specific primers (see Table 1). qPCR was performed using Low ROX SYBR Master Mix dTTP Blue (Takyon, #UF-LSMT_B0701) on an Applied Biosystems StepOnePlus Real-Time PCR System. Data were analyzed with the ΔCt method.

**Bulk RNA-sequencing and analysis** 10 guts of 7 days post-fertilization WT and *il22*$^{-/-}$ larvae conventionally raised, germ-free or co-housed were dissected per replicates (4 replicates per group). Total RNA was extracted with the Single-Cell RNA Purification kit (Norgen Biotek, #51800) following manufacturer's instructions. The RNA integrity and concentration were analyzed on Agilent 4200 Tapestation system using the high sensitivity RNA ScreenTape Analysis kit (Agilent, #5067-5579) and apparatus. RNA sequencing libraries were prepared from 500 ng of total RNA using the Illumina TruSeq Stranded mRNA Library preparation kit. cDNA quality was checked on Agilent 2100 Bioanalyzer using Agilent High Sensitivity DNA kit (Agilent #5067-4626). After quality control, libraries were sequenced with 100-bp paired-end (PE100) reads on the NovaSeq 6000 (Illumina) sequencer. Raw data were checked for quality using FastQC (v0.11.8) and aligned to the reference genome for Danio rerio danRer11 from Genome Reference Consortium. Analysis strategy includes unsupervised analyses such as PCAs and differential expression analyses (done with DEBrowser bioconductor package or on R with edgeR package).

**Microbial DNA extraction and 16S rRNA sequencing** Dissection of 15 intestines from WT and *il22*$^{-/-}$ 7 dpf larvae per replicates followed by bacterial DNA isolation using the DNeasy PowerSoil kit (Qiagen, #47014) following manufacturer's instructions. They were directly stored at -20°C until sequencing. Two primers were used to amplify the 16S rRNA genes covering the hypervariable regions V3 to V4: 338F: ACTCCTACGGGAGGCAGCAG and 806R: GGACTACH-VGGGTWTCTAAT. Amplified regions were sequenced by BGI technologies using the DNBSEQ™ sequencing technology platform.

From the resulting raw data, redundancy analysis was done with Canoco 5.15 (*43*) with ASV relative abundance as response variables, after transforming with the formula log(1000*relative_abundance + 1). RDA p-values were determined through permutation testing (500 permutations). Bacterial genome sequences were downloaded with the NCBI tool "datasets" (https://www.ncbi.nlm.nih.gov/datasets/docs/v2/reference-docs/command-line/datasets/), 16S sequences were extracted with BioPython (https://biopython.org/). ASVs were compared to 16S sequences with blastn 2.9 (*44*). HMM screening was done with hmmsearch from HMMER3.3 (http://hmmer.org).

**Generation of germ-free (GF) larva and co-housing experiments** Fertilized zebrafish eggs were treated with bleach (0,05% active chloride) for maximum 2 min at 3–4 hpf and then washed twice with sterile E3 medium for 5 min. Fertilized eggs were incubated in chlorine hypochlorite (0.003% active chloride) for 20 minutes. After washing, embryos were left in sterile E3 medium containing Ampicillin (200 µg/mL), Kanamycin (5 µg/mL), Ceftazidime (200ug/mL) and Chloramphenicol (20ug/mL) and placed at 28 °C in isolated containers. Media was renewed every day in sterile conditions until the day of sample collection. Sterility of larvae and E3 water was monitored every 2 days by incubating fish water in TBS media for 24h at 37°C. GF zebrafish were co-housed from 4 dpf to 7 dpf with conventionally-raised larvae placed in a 40um filter (Fisherbrand, #22-363-547) to allow microbiota transfer. At 7 dpf, samples were processed for RT-qPCR or further analyzed by live imaging.

**Recording *in vivo* intestinal motility** 7 dpf larvae were anesthetized using 100 ug/ml of Tricaine for several minutes at 28°C. Larvae were then embedded in a liquified 0,5% low melting point agarose (Promega, #V2111) and covered with E3 water containing tricaine (100 ug/ml). Imaging was performed using the THUNDER Imager Model Organism (Leica). Movies were taken with 100ms exposure time at 10 frames per second. They were later analyzed by making kymographs on 180um length region of the larval intestine using the "Velocity Measurement Tool" macro (https://dev.mri.cnrs.fr/projects/imagej-macros/wiki/Velocity_Measurement_Tool) in ImageJ software (NIH).

**Food transit experiment** Larvae were trained from 4 dpf to 7 dpf with usual food. On test day, larvae were fed during 2h with food coupled with non-digestive fluoresphere carboxylate 2um (Invitrogen, #F8827). Only fish having the intestinal bulb filled with fluorescent food were used for the experiment. Pictures were taken using the THUNDER Imager Model Organism (Leica) 3, 6, 12 and 24h after feeding. The distance between the anterior end of the intestinal bulb and the fluorescent food in the intestine was measured using ImageJ software (NIH).

**Zebrafish *Edwarsiella tarda* infection** The day before challenging *Edwarsiella tarda* FL60 (kindly provided by Dr. Phillip Klesius (USDA, Agricultural Research Service, Aquatic Animal Health Research Unit), the bacteria grew in TSB medium + tetracycline (15ug/mL) at 28ºC overnight. On the day of the challenge, a 1:100 dilution was performed, and the bacteria grew to OD600 = 0.250 (approximately 108 CFU/mL). The bacteria were then centrifuged twice at 4500 rpm for 5-10 minutes and resuspended in E3 1X water to OD600 = 0.250. Six larvae per 6 ml of liquid were incubated in E3 water containing bacteria for 5 hours at 28°C. The infected larvae undergo three washes with E3 1X water. Survival was monitored every 12 hours for 3 days post infection.

**Chemical treatments in larval zebrafish** *Gut injection*: 6 to 7 dpf larvae were microinjected in the gut with Indole (Sigma, #I3408). 4h post-injection, larval intestines were dissected and stored at -20°C or directly processed for RT-qPCR.

*Water incubation*: A groups of 25 larvae were kept in E3 1X medium (controls) or in water coming from WT larvae co-housed *il22*$^{-/-}$ larvae. The medium was changed daily. The experiment was done at least 3 times with different egg batches.
*Incubation with chemical compounds:* 1. 7 dpf larvae were incubated with DMSO or Trpa1 receptor inhibitor (Sigma, #HC030031) at 10uM for 4h before dissection for qPCR. 2. Larvae were also incubated with PBS 1X or ghrelin 100nM (Genscript, # RP10781) from 4 dpf to 7 dpf. Freshly prepared ghrelin peptide was added daily and live imaging was performed on treated samples at 7 dpf. 3. WT CV larvae co-housed with *il22*$^{-/-}$ GF were incubated with DMSO or ghrelin receptor antagonist at 50ng/uL (Phoenix pharmaceuticals, #031-22) for 24h before imaging for gut motility measurements.

**Food transit experiment in mice** WT and *Il22*$^{-/-}$ mice male or female aged of 3-4, 5 or 8-10 weeks were used. Carmine red (Sigma, #C6152) was given by gavage to 3h-fasted mice (10 mg/ml of water, 10 µl/g body). The total intestinal transit time was measured by determination of time between ingestion of carmine red and first appearance of the dye in the feces.

**Statistical analysis** Statistical analyses were conducted using Rstudio or GraphPad Prism. The types of statistical tests and significance levels are described in respective figure legends. The results were considered statistically significant when P value was lower than 0.05 and were marked in the figures as **** $P< 0.0001$, ***$P < 0.001$, **$P < 0.01$, and *$P < 0.05$.

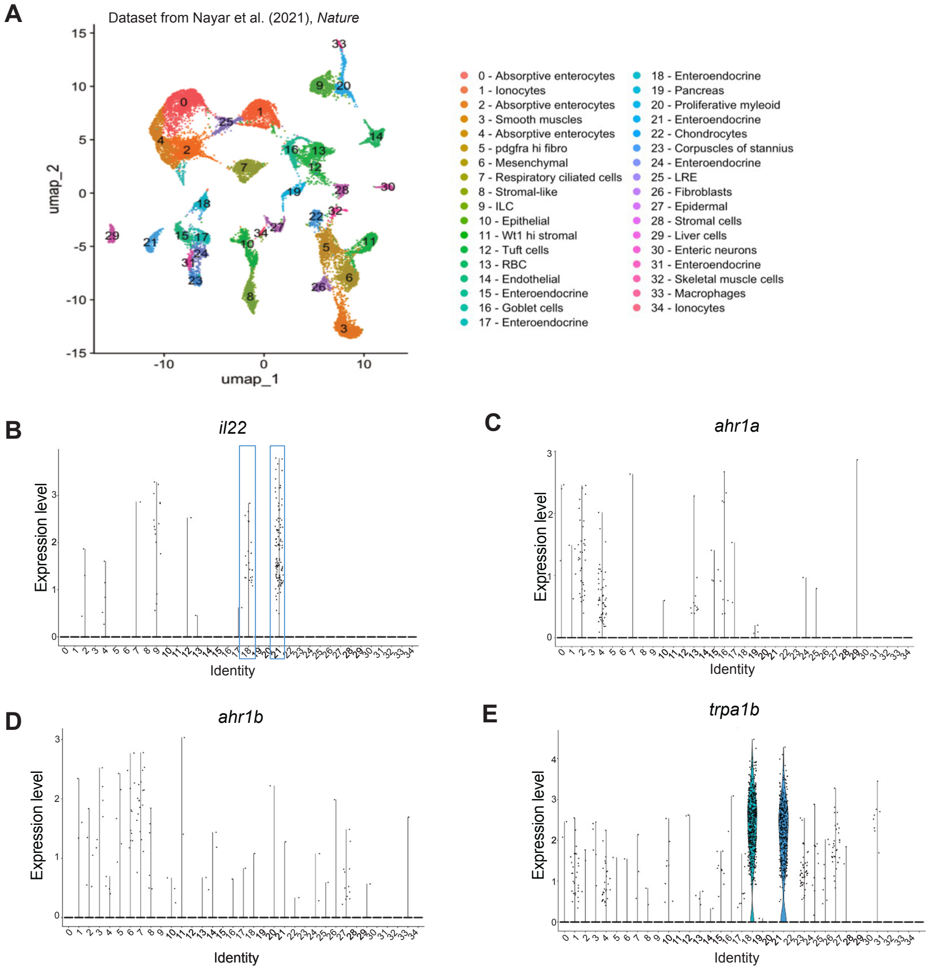


**Fig S1. Re-analysis of scRNAseq dataset of zebrafish larval gut (Nayar et al., Nature 2021).** **A.** Re-analysis of scRNA-seq of larval intestines, uniform manifold approximation and projection (UMAP) visualization of cell type clusters. **B, C, D, E**. Violin plots for the expression level of the indicated genes. Blue boxes on **(B)** are highlighting *trpa1b*$^{+}$ enteroendocrine cells clusters.

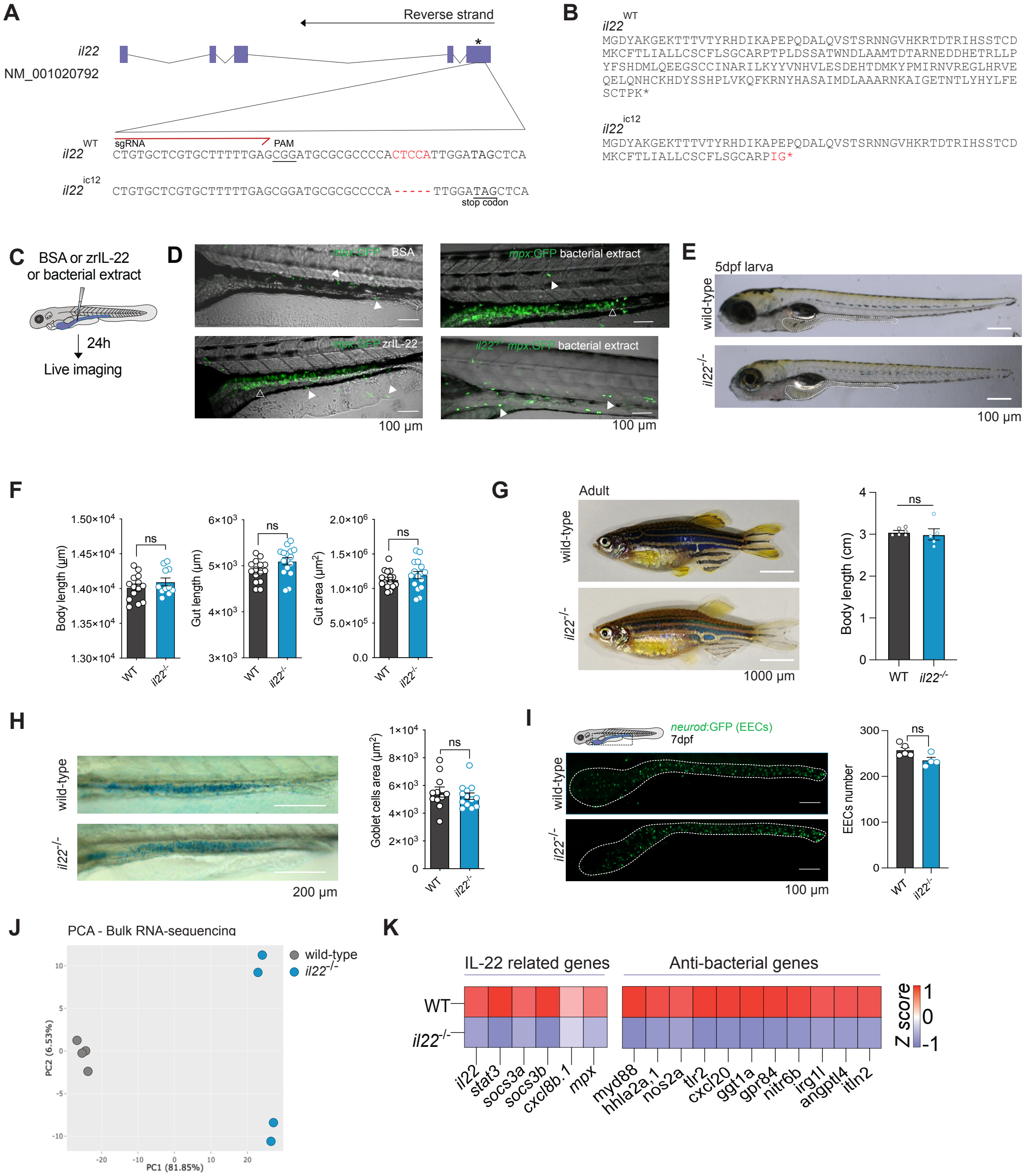
A
Reverse strand
il22
NM_001020792
*
il22 WT
sgRNA
PAM
CTGTGCTCGTGCTTTTTGAGCGGATGCGCGCCCCACTCCATTGGATAGCTCA
il22 ic12
CTGTGCTCGTGCTTTTTGAGCGGATGCGCGCCCCA-----TTGGATAGCTCA
stop codon
B
il22 WT
MGDYAKGEKTTTVTYRHDIKAPEPQDALQVSTSRNNGVHKRTDTRIHSSTCD
MKCFTLIALLCSCFLSGCARPTPLDSSATWNDLAAMTDTARNEDDHETRLLP
YFSHDMLQEEGSCCINARILKYYVNHVLESDEHTDMKYPMIRNVREGLHRVE
QELQNHCKHDYSSHPLVKQFKRNYHASAIMDLAAARNKAIGETNTLYHYLFE
SCTPK*
il22 ic12
MGDYAKGEKTTTVTYRHDIKAPEPQDALQVSTSRNNGVHKRTDTRIHSSTCD
MKCFTLIALLCSCFLSGCARPIG*
C
BSA or zrIL-22
or bacterial extract
24h
Live imaging
D
mpx:GFP BSA
mpx:GFP bacterial extract
mpx:GFP zrIL-22
il22-/- mpx:GFP bacterial extract
100 µm
100 µm
E
5dpf larva
wild-type
il22-/-
100 µm
F
Body length (µm)
Gut length (µm)
Gut area (µm²)
ns
ns
ns
WT
il22-/-
G
Adult
wild-type
il22-/-
1000 µm
Body length (cm)
ns
WT
il22-/-
H
wild-type
il22-/-
200 µm
Goblet cells area (µm²)
ns
WT
il22-/-
I
neurod:GFP (EECs)
7dpf
wild-type
il22-/-
100 µm
EECs number
ns
WT
il22-/-
J
PCA - Bulk RNA-sequencing
wild-type
il22-/-
PC1 (81.85%)
PC2 (6.53%)
K
IL-22 related genes
Anti-bacterial genes
WT
il22-/-
Z score
il22
stat3
socs3a
socs3b
cxcl8b.1
mpx
myd88
hhla2a.1
nos2a
tlr2
cxcl20
ggt1a
gpr84
nitr6b
irg1l
angptl4
itln2

**Fig S2**. **Characterization of the *il22*$^{-/-}$ zebrafish line**

**A.** Schematic for the mutation generated in the zebrafish *il22* gene (-5bp) by CRISPR/Cas9**. B.** Predicted protein sequence for Il22 in wild-type (WT) and *il22* mutant individuals, according to the DNA sequences obtained. **C, D.** Validation of our new *il22*$^{-/-}$ line. To confirm the non-functionality of the *il22* gene in our new mutant line, we initially sought to identify an IL-22-dependent gene. Prior studies using the Tg(*mpx*:GFP) line showed that the myeloid-specific peroxidase (*mpx*) gene, typically expressed in neutrophils, is highly induced in intestinal epithelial cells (IECs) following injection of zebrafish recombinant IL-22 (zrIL-22 (*45*)). We corroborated this finding by injecting BSA (D top left panel) or zebrafish recombinant IL-22 (zrIL-22) protein (D bottom left panel) in guts of 5 dpf Tg(*mpx*:GFP) larvae. Subsequently, we microinjected a bacterial extract known to induce *il22* expression in zebrafish guts in Tg(*mpx*:GFP) (D top right panel) and *il22*$^{-/-}$Tg(*mpx*:GFP) (D bottom right panel). The absence of GFP induction in IECs after bacterial extract injection in the *il22*$^{-/-}$Tg(*mpx*:GFP) larvae confirmed that the *il22* gene in our newly generated line is non-functional. Filled white arrowheads show neutrophils and empty arrowheads highlight epithelial cells. Scale bar = 100 um. **E.** Representative images of 5 dpf WT and *il22*$^{-/-}$ larvae. Dotted lines enclose the gut. Scale bar = 100 um. **F.** Quantification of body length, gut length, and gut area in 5 dpf WT and *il22*$^{-/-}$ larvae. **G.** Brightfield pictures of adult WT and *il22*$^{-/-}$ fish and quantification of their body length. Scale bar = 1000 um. **H.** Alcian blue staining of 5 dpf WT and *il22*$^{-/-}$ and quantification of the stained area in the gut. **I.** Confocal microscopy of Tg(*neurod*:GFP), labelling EECs, in WT or *il22*$^{-/-}$ 7 dpf larvae and quantification of the number of GFP positive cells in the gut. Dotted lines surround the gut. Scale bar = 100 um. **J.** PCA plot of bulk RNA-sequencing comparing 7 dpf WT and *il22*$^{-/-}$ larval intestines. **K.** Heatmap showing expression levels of the indicated IL-22-regulated genes in WT and *il22*$^{-/-}$ guts. In mammals, IL-22 activates the STAT3 signaling pathway and anti-bacterial peptide production in IECs (*46*). *il22*$^{-/-}$ larval guts showed a significant decrease in the expression of *il22*, as well of *stat3*, *socs3a/b,* and *mpx*, a known anti-bacterial gene (*45*). Statistical analysis was performed with Mann-Whitney * $P$ < 0,05, ** $P$ < 0,01. Data are representative of at least two independent experiments.

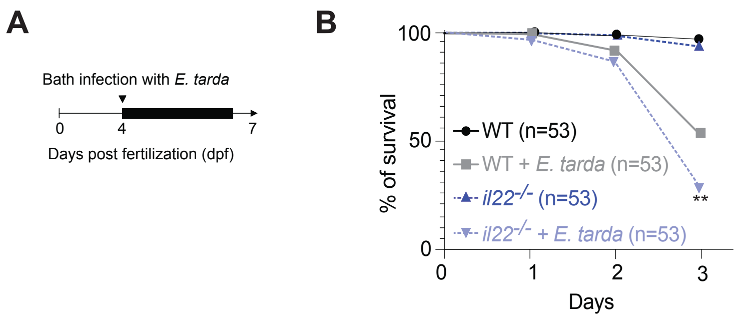


**Fig S3. Zebrafish IL-22 protects from gut bacterial infection in larvae**
**A.** Experimental strategy for a bath infection with live *Edwardsiella tarda* of WT and *il22*$^{-/-}$ larvae. Larvae were infected from 4 to 7 dpf and survival was measured daily. **B.** Survival curves of WT and *il22*$^{-/-}$ with or without *E. tarda* infection. Statistical analysis was performed with log-rank test ** $P < 0{,}01$. Data are representative of at least two independent experiments.

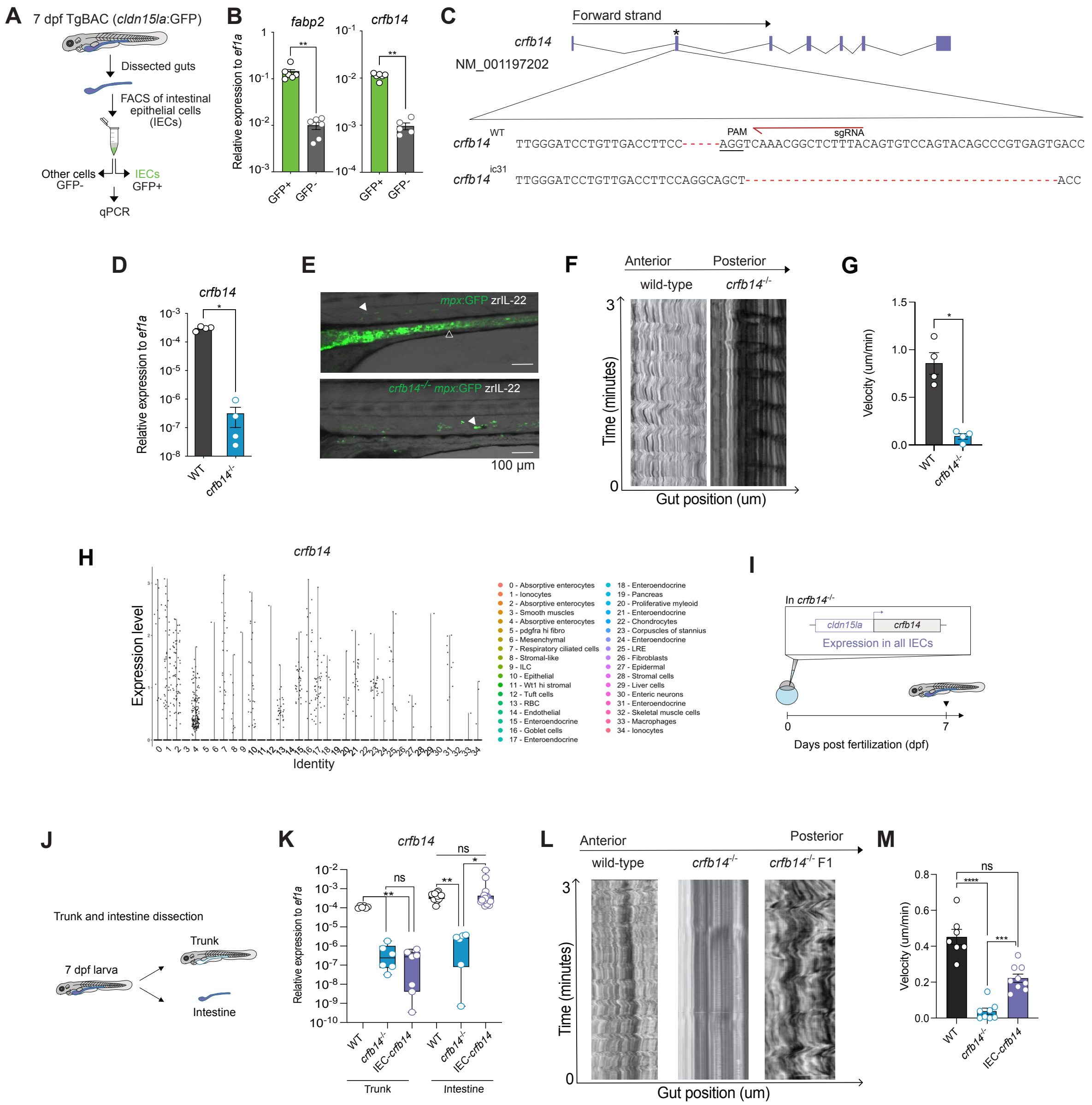
A
7 dpf TgBAC (cldn15la:GFP)
Dissected guts
FACS of intestinal epithelial cells (IECs)
Other cells GFP-
IECs GFP+
qPCR
B
fabp2
crfb14
Relative expression to ef1a
C
Forward strand
crfb14
NM_001197202
PAM
sgRNA
crfb14 WT TTGGGATCCTGTTGACCTTCC----AGGTCAAACGGCTCTTTACAGTGTCCAGTACAGCCCGTGAGTGACC
crfb14 ic31 TTGGGATCCTGTTGACCTTCCAGGCAGCT------------------------------------------ACC
D
E
mpx:GFP zrIL-22
crfb14-/- mpx:GFP zrIL-22
100 μm
F
Anterior
Posterior
wild-type
Time (minutes)
Gut position (um)
G
Velocity (um/min)
H
Expression level
Identity
0 - Absorptive enterocytes
1 - Ionocytes
2 - Absorptive enterocytes
3 - Smooth muscles
4 - Absorptive enterocytes
5 - pdgfra hi fibro
6 - Mesenchymal
7 - Respiratory ciliated cells
8 - Stromal-like
9 - ILC
10 - Epithelial
11 - Wt1 hi stromal
12 - Tuft cells
13 - RBC
14 - Endothelial
15 - Enteroendocrine
16 - Goblet cells
17 - Enteroendocrine
18 - Enteroendocrine
19 - Pancreas
20 - Proliferative myeloid
21 - Enteroendocrine
22 - Chondrocytes
23 - Corpuscles of stannius
24 - Enteroendocrine
25 - LRE
26 - Fibroblasts
27 - Epidermal
28 - Stromal cells
29 - Liver cells
30 - Enteric neurons
31 - Enteroendocrine
32 - Skeletal muscle cells
33 - Macrophages
34 - Ionocytes
I
In crfb14-/-
cldn15la
crfb14
Expression in all IECs
Days post fertilization (dpf)
J
Trunk and intestine dissection
7 dpf larva
Trunk
Intestine
K
IEC-crfb14
L
crfb14-/- F1
M

**Fig S4. Expression of the IL-22 receptor specifically in gut epithelial cells is sufficient to sustain gut motility zebrafish**

**A**. Schematic representation of FACS sorting of *cldn15la*:GFP-positive and -negative cells from dissected larval guts, followed by RT-qPCR analysis. **B.** RTqPCR analysis measuring the expression of *fabp2* (epithelial marker) and *crfb14* in sorted cells. **C.** Schematic of the mutation generated in the zebrafish *crfb14* gene by CRISPR/Cas9, leading to a 5 bp insertion and a 43 bp deletion in exon 2, which removes an essential splicing site. **D.** RT-qPCR analysis on dissected guts measuring *crfb14* gene expression in WT and the new *crfb14*$^{-/-}$ line. *crfb14*$^{-/-}$ had significantly reduced *crfb14* expression in the gut. **E.** Representative images of 5 dpf Tg(*mpx*:GFP) larvae (top), and *crfb14*$^{-/-}$ Tg(*mpx*:GFP) larvae (bottom) injected with zrIL-22. Filled white arrowheads show neutrophils and empty arrowheads highlight IECs. Scale bar = 100 um. zrIL-22 microinjection into *crfb14*$^{-/-}$ larvae failed to induce IL-22-target gene expression, confirming *crfb14* as the zebrafish IL-22-specific receptor chain. **F**. Representative kymograph analysis of 180 um in the midgut of 7 dpf WT and *crfb14*$^{-/-}$ larvae using ImageJ. **G.** Gut motility speed in 7 dpf WT and *crfb14*$^{-/-}$ larvae. **H.** Re-analysis of a single-cell RNA-sequencing dataset of zebrafish larval guts (Nayar et al, 2020). **I.** Schematic of the strategy utilized to generate a zebrafish line expressing *crfb14* only in IECs. Expression of *crfb14* is driven by an IEC-specific promoter (*cldn15la*) in a *crfb14*$^{-/-}$ background (IEC-*crfb14* line). **J.** Schematic of the trunk and intestinal tissue collection of 7 dpf WT and *crfb14*$^{-/-}$ larvae for RNA extraction. **K.** RT-qPCR for *crfb14* in dissected trunks or guts of 7 dpf WT, *crfb14*$^{-/-}$ and IEC-*crfb14* larvae. The IEC-*crfb14* line showed *crfb14* expression only in the gut. **L.** Examples of kymograph analysis of 180 um in the midgut of 7 dpf WT, *crfb14*$^{-/-}$ and IEC-*crfb14* (*cldn15la*:*crfb14* line) larvae using ImageJ. **M.** Gut motility speed in 7 dpf WT, *crfb14*$^{-/-}$ and IEC-*crfb14* larvae**.** Statistical analysis was performed with Mann-Whitney : ns: not significant, * $P < 0,05$, ** $P < 0,01$. These data are representative of at least three independent experiments.

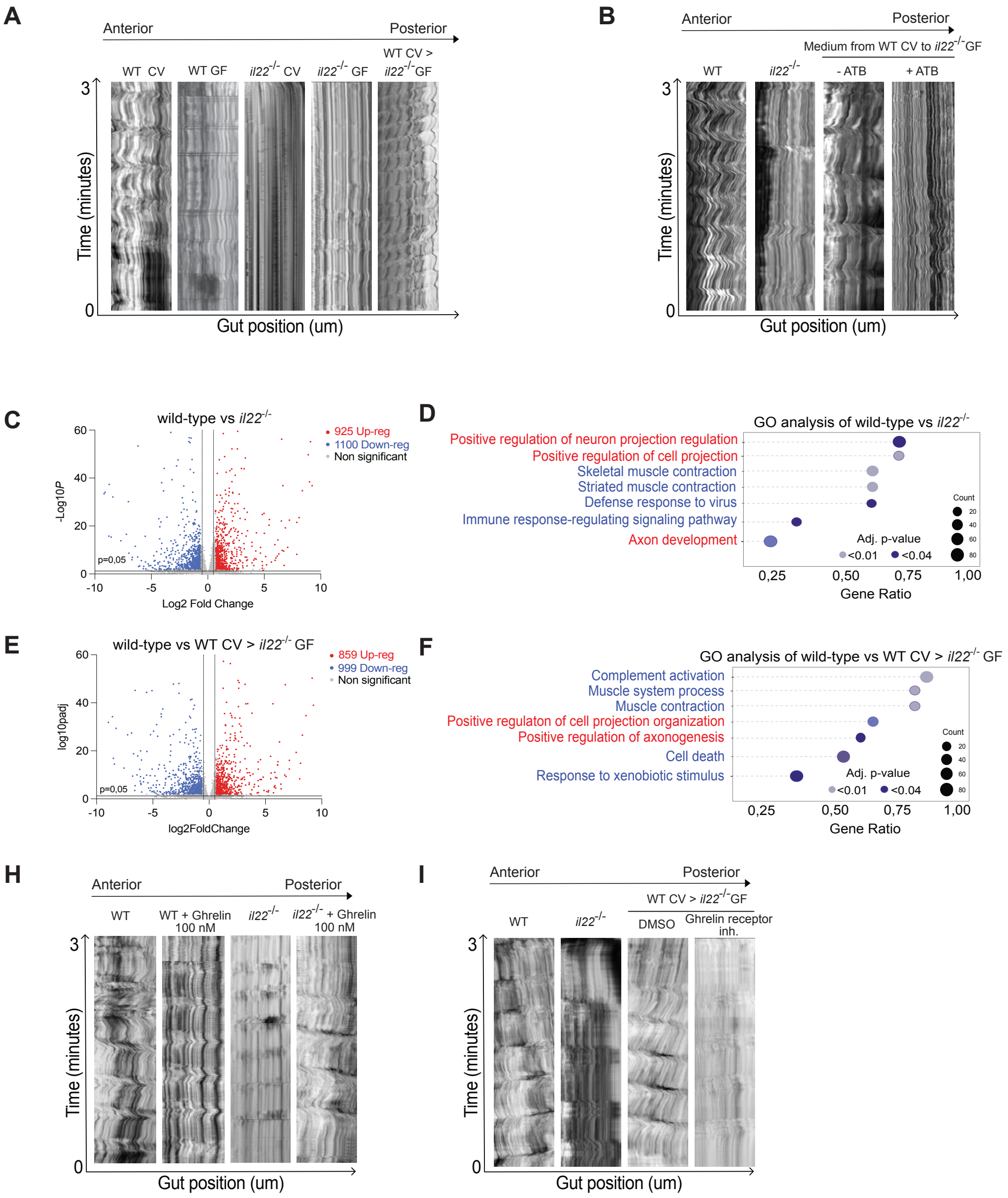
A
Anterior
Posterior
WT CV
WT GF
il22-/- CV
il22-/- GF
WT CV > il22-/- GF
Time (minutes)
3
0
Gut position (um)
B
Anterior
Posterior
Medium from WT CV to il22-/- GF
WT
il22-/-
- ATB
+ ATB
Time (minutes)
3
0
Gut position (um)
C
wild-type vs il22-/-
925 Up-reg
1100 Down-reg
Non significant
-Log10P
p=0,05
Log2 Fold Change
D
GO analysis of wild-type vs il22-/-
Positive regulation of neuron projection regulation
Positive regulation of cell projection
Skeletal muscle contraction
Striated muscle contraction
Defense response to virus
Immune response-regulating signaling pathway
Axon development
Count
Adj. p-value
<0.01
<0.04
0,25
0,50
0,75
1,00
Gene Ratio
E
wild-type vs WT CV > il22-/- GF
859 Up-reg
999 Down-reg
Non significant
log10padj
p=0,05
log2FoldChange
F
GO analysis of wild-type vs WT CV > il22-/- GF
Complement activation
Muscle system process
Muscle contraction
Positive regulaton of cell projection organization
Positive regulation of axonogenesis
Cell death
Response to xenobiotic stimulus
Count
Adj. p-value
<0.01
<0.04
0,25
0,50
0,75
1,00
Gene Ratio
H
Anterior
Posterior
WT
WT + Ghrelin 100 nM
il22-/-
il22-/- + Ghrelin 100 nM
Time (minutes)
3
0
Gut position (um)
I
Anterior
Posterior
WT CV > il22-/- GF
WT
il22-/-
DMSO
Ghrelin receptor inh.
Time (minutes)
3
0
Gut position (um)

**Figure S5. Transcriptomic analysis of guts of *il22*$^{-/-}$ co-housed with WT larvae**
**A.** Examples of kymograph analysis of 180 um in the midgut of 7 dpf WT CV, WT GF, *il22*$^{-/-}$ CV, *il22*$^{-/-}$ GF, or *il22*$^{-/-}$ GF co-housed with WT CV. **B.** Examples of kymograph analysis of 180 um in the midgut of 7 dpf *il22*$^{-/-}$ GF larvae after transferring the medium from WT CV either treated or not with antibiotics (see Material and methods) **C**. Volcano plot showing the dysregulated genes in guts of *il22*$^{-/-}$ compared to WT larvae with a $P < 0,05$ and a fold change > 1.5. Downregulated genes are labeled in blue and upregulated genes are shown in red. **D.** Gene ontology analysis of differentially expressed genes in WT versus *il22*$^{-/-}$. GO terms for upregulated genes are shown in red and GO terms for downregulated genes are shown in blue. **E**. Volcano plot showing the dysregulated genes in guts of WT CV compared to *il22*$^{-/-}$ GF co-housed with WT CV larvae with a $P < 0,05$ and a fold change > 1.5. Downregulated genes are labeled in blue and upregulated genes are shown in red. **F.** Gene ontology analysis of differentially expressed genes in WT versus to *il22*$^{-/-}$ GF co-housed with WT CV. GO terms for upregulated genes are shown in red and GO terms for downregulated genes are shown in blue. **G**. Examples of kymograph analysis of 180 um in the midgut of 7 dpf WT, *il22*$^{-/-}$ and *il22*$^{-/-}$ incubated or not with Ghrelin protein. **H**. Representative kymograph analysis of 180 um in the midgut of 7 dpf of WT CV, *il22*$^{-/-}$ CV, *il22*$^{-/-}$ GF co-housed with WT CV larvae treated with DMSO or with the Ghrelin receptor inhibitor. These data are representative of at least three independent experiments.

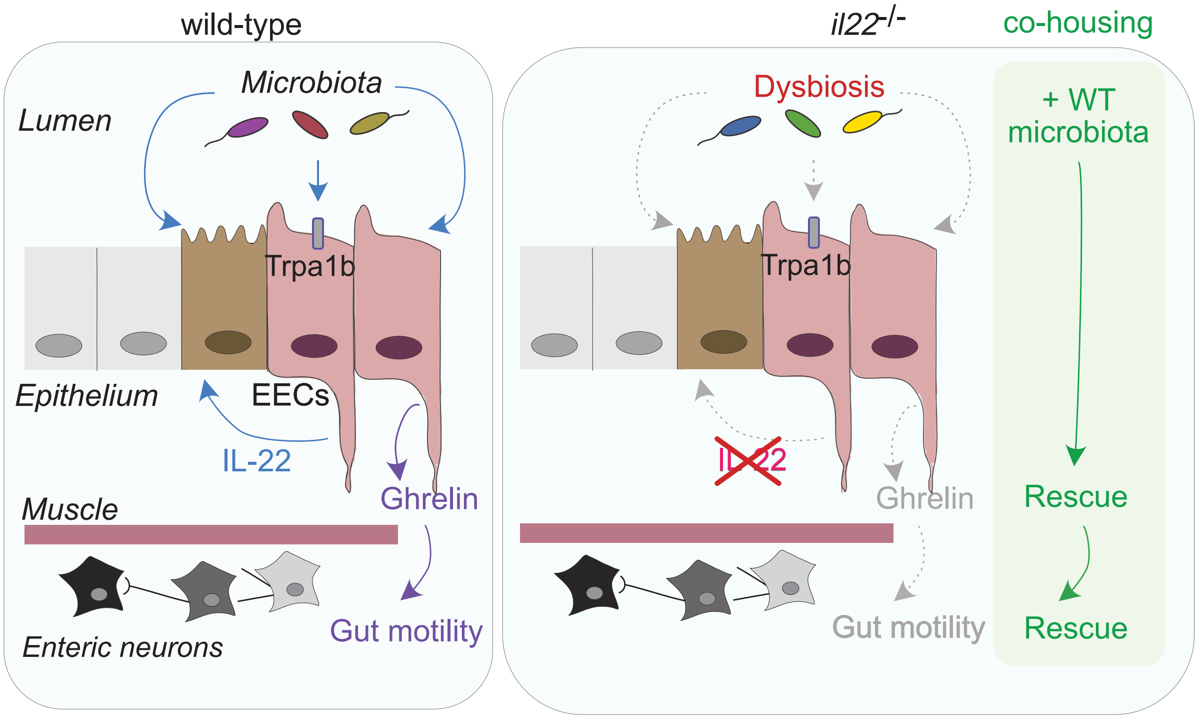


**Fig S6. Proposed model for the role of IL-22 in controlling gut motility during early life**

In this study, we discovered that IL-22 is produced by enteroendocrine cells in larvae and plays a critical role in gut immunity and hormone expression. IL-22 expression is regulated by the microbiota, specifically by tryptophan metabolites that signal through the Trpa1 receptor. In parallel, we found that IL-22 is involved in controlling the composition of the microbiota in zebrafish larvae, similar to its role in mammals. Interestingly, we observed defects in neuronal activity and intestinal motility in IL-22-deficient larvae. Transfer of bacteria from wild-type zebrafish to *il22*$^{-/-}$ larvae restored gut motility by restoring ghrelin hormone expression, a key regulator of intestinal motility. These findings highlight a circuit in which IL-22 plays a critical role in the maintenance of the microbiota and the hormonal balance during early development in order to ensure proper gut physiology.

| RESSOURCE | SOURCE | IDENTIFIER |
|---|---|---|
| **Antibodies** | | |
| Mouse anti-HuC/D monoclonal antibody | Invitrogen | Cat# A21271 |
| Rabbit anti-Ghrelin monoclonal Antibody | Abcam | ab209790 |
| Chicken anti-GFP Polyclonal antibody | Abcam | ab13970 |
| Living Colors anti DsRed Polyclonal Antibody | TAKARA | Cat# 632496, RRID:AB_10013483 |
| Goat anti-Chicken AF488 | Life technologies | Cat# A-11039 |
| Goat anti-Rabbit Cy3 | Jackson Immunoresearch | *Cat# 111-166-003* |
| Goat anti-Mouse AF647 | Life technologies | Cat# A32728 |
| **Oligonucleotides** | | |
| RTqPCR for zebrafish gene: *il22* | Eurofins Genomics | TGCAGAATCACTGTAAACACGA<br>CTCCCCGATTGCTTTGTTAC |
| RTqPCR for zebrafish gene: *crfb14* | Eurofins Genomics | AACGGCTCTTTACAGTGTCCA<br>TGCATCCATCACATCAGTCAGA |
| RTqPCR for zebrafish gene: *fabp2* | Eurofins Genomics | TGGGCGTCACCTTTGACTAT<br>GCGTGTCTCCCTCTATGACC |
| RTqPCR for Mouse gene: *Ghrl* | Eurofins Genomics | GAAGCCACCAGCTAAACTGCAG<br>CTGACAGCTTGATGCCAACATCG |

| | | |
|---|---|---|
| **KASP assay** | | |
| Primers for the zebrafish gene : *il22* | LGC Biosearch Technologies | Primer allele X GGTGGCTGAGCTATCCAATGGA<br>Primer allele Y GTGGCTGAGCTATCCAATGGG<br>Primer common CTTATTGCTTTGCTGTGCTCGTGCTT |

**Table S1. Antibodies and primers sequences**

| Larvae | Number of *il22*:mCherry⁺ cells | Number of *il22*:mCherry⁺ *neurod*:GFP⁺ cells | Total number of *il22*:mCherry cells | Pourcentage of double positive cells (%) |
|---|---|---|---|---|
| 1 | 0 | 2 | 2 | 100 |
| 2 | 0 | 1 | 1 | 100 |
| 3 | 0 | 2 | 2 | 100 |
| 4 | 1 | 2 | 3 | 66,7 |
| 5 | 0 | 2 | 2 | 100 |
| 6 | 0 | 4 | 4 | 100 |
| 7 | 2 | 4 | 6 | 66,7 |

**Table S2. Most intestinal epithelial cells expressing *il22* are enteroendocrine cells**
